# Biofabrication of a filtration barrier by integrating electrospun membranes and flow in a glomerular co-culture

**DOI:** 10.1101/2024.12.05.626949

**Authors:** Camilla Mussoni, Anna Rederer, Vladimir Stepanenko, Frank Würthner, Philipp Stahlhut, Jürgen Groll, Mario Schiffer, Taufiq Ahmad, Janina Müller-Deile

**Affiliations:** Department of Functional Materials in Medicine and Dentistry, Institute of Functional Materials and Biofabrication (IFB), and Bavarian Polymer Institute (BPI), Julius-Maximilians-Universität Würzburg, Würzburg, Germany; Department of Nephrology, Medizinische Klinik 4, Universitätsklinium Erlangen, Friedrich-Alexander-Universität Erlangen-Nürnberg; Institut für Organische Chemie & Center for Nanosystems Chemistry (CNC), Julius-Maximilians-Universität Würzburg, Würzburg, Germany

## Abstract

The glomerulus is the functional unit of the kidney, where urine is filtered from blood. This process happens through the glomerular filtration barrier (GFB) which is composed of glomerular endothelial cells, podocytes, and glomerular basement membrane (GBM). Damage to any component of GFB results in failure of the barrier’s function, causing proteinuria that can lead to end-stage kidney failure. There is a high need for reliable *in vitro* models of the GFB to study pathological conditions and to test potential novel therapeutic options. We established an artificial GBM generated by electrospinning of poly-L-lactic acid fibers that were coated with polydopamine and gelatin and seeded with human glomerular endothelial cells and podocytes onto different sides. The orientation of fibers in the artificial GBM, their surface chemistry and effects on glomerular cells were characterized in depth. Glomerular endothelial cells and podocytes, including hiPSC-derived podocytes formed monolayers on the artificial GBM and revealed cell type-specific marker expression and morphology. Cell-cell communication was possible between podocytes and glomerular endothelial cells in both directions though the artificial membrane. Different molecular dextrans showed size selective permeability of the *ex vivo* GFB model. Introduction of shear stress by applying flow with a 3D printed micro-bioreactor improved cellular ultrastructure with formation of glomerular endothelial cell fenestrae-like structures and long podocyte foot process-like protrusions that both are usually absent in other *in vitro* models. Personalized hiPSC-derived podocytes within our model will allow to study the role of patient-specific podocyte mutations or individual treatment response *ex vivo* in the future.

## INTRODUCTION

Kidney glomeruli serve as the primary site for selective ultrafiltration of blood involved in the excretion of metabolic waste by allowing water and small molecules to pass while preventing the passage of larger proteins and blood cells [1]. The glomerular filtration barrier (GFB) is a tri-layered structure composed of fenestrated endothelium with 70–100 nm diameter fenestrations, a glomerular basement membrane (GBM) rich in extracellular matrix (ECM) proteins such as type IV collagen and laminin and podocytes with interdigitating foot processes connected by slit diaphragms [1b, 2]. This architecture ensures size and charge selectivity and enables the daily filtration of approximately 180 liters of primary urine from whole blood [1]. Any disruption in the component of the glomerulus could lead to proteinuria, ultimately leading to kidney failure [3]. Therefore, proper development, regulation, and cross-communication between all layers of the GFB is essential for the maintenance and functionality of the barrier. Different glomerular diseases like diabetic nephropathy or primary glomerulonephropathies caused by autoimmune dysregulation or auto-antibodies against components of the GFB but also genetic diseases due to mutations in podocyte genes or genes encoding proteins of the GBM lead to impairments in GFB function with proteinuria and/or hematuria and progressive chronic kidney disease (CKD).

There is a huge need for *ex vivo* models of the GFB to simulate physiological and pathophysiological conditions. However, conventional two-dimensional (2D) culture methods are limited in replicating complex physiological and microarchitectures of GFB. Therefore, there is an increasing demand for *in vitro* models that can accurately simulate normal and pathological conditions. Such models need to retain critical features of the GFB, including the fenestration of glomerular endothelial cells, the interdigitating foot processes of podocytes, and the ability to perform size- and charge-selective filtration.

The advancement of GFB *in vitro* models has been driven by the integration of 3D culture systems, organ-on-chip (OOC) platforms [4], and biomimetic materials such as nanofibers and hydrogels [4g, 4j, 5], which enhance cell support, signaling, and morphogenesis. OOC platforms, capable of mimicking the complex dynamic environment of the glomerulus, have emerged as a transformative tool in preclinical research. Despite these advances, traditional OOC systems rely heavily on synthetic membranes like polydimethylsiloxane (PDMS) and polycarbonate (PC), which are significantly thicker than human basement membranes, limiting physiological accuracy. These synthetic materials are non-degradable, flat, and lack the necessary topographical features to support proper ECM remodeling and transmembrane crosstalk, both essential for studying tissue functions and disease mechanisms. Nanofiber scaffolds provide a promising alternative to PDMS and PC membranes by offering ultrathin, biomimetic substrates with high surface area and tunable porosity that better replicate native tissue structures [6].

Here, we developed an artificial GFB by using an electrospun poly-L-lactic acid (PLLA) membrane to mimic the GBM that was seeded with human glomerular endothelial cells and human podocytes (immortalized or hiPSC-derived). Electrospun nanofibers offer numerous advantages over the commercially available synthetic systems in terms of size, shape, variety of materials, fiber morphology, and orientations, and the modification of nanofibers with bioactive cues. By fabricating the artificial GBM in different orientations, we demonstrated that cell attachment and morphology are impacted by fiber topography. Nevertheless, we could achieve high viable cell monolayers with characteristic glomerular cell morphology and marker expression on aligned membrane. Functional properties of the GFB model, such as barrier function, were investigated by measuring dextran passage through the filtration barrier. Moreover, we designed and fabricated with 3D printing a customized reusable bioreactor to apply flow conditions for five days. We observed fenestrae-like pores organized in clusters on the glomerular endothelial cell surface confirming the maturation of cells within the system. Furthermore, we were able to demonstrate glomerular cell-cell communication within our model and selective permeability.

Thus, we present an *in vitro* model of the GFB mimicking the barrier on ultrastructure with glomerular endothelial cell fenestrae-like and podocyte foot processes-like, allowing for flow condition and to measure glomerular barrier function and glomerular permeability in healthy and diseased condition. Using personalized hiPSC-derived podocytes in this model allows for individualized disease model in the future.

## MATERIALS AND METHODS)

### Experimental Design

#### Generation of electrospun membranes and their surface modification

We fabricated the nanofibers mesh using electrospinning to artificially mimic the glomerular basement membrane. A 2% solution of poly-L-lactic acid (PLLA) (Purasorb® PL65, Corbion, Amsterdam, Netherlands) was prepared in HexaFluoroIsoPropanol (HFIP) (Sigma Aldrich, Germany). Then, the prepared solution was extruded at the rate of 1 mL/h using a syringe pump (World Precision Instruments, Sarasota, Florida, USA) through a 27G nozzle under the voltage of 13 kV applied to the nozzle. The nanofibers were collected on the aluminum foil covered on the rotating grounded mandrel (Ø 10 cm), positioned 15 cm from the nozzle. The random nanofibers were collected on the mandrel with a rotating speed of 180 rpm. The collector setup was modified to achieve aligned nanofibers by replacing the aluminum foil with a Teflon® film decorated with copper microwires spaced 1 cm apart and wrapped around the grounded mandrel, rotating at 500 rpm to collect aligned nanofibers.

For surface coating with polydopamine, membranes were dip-coated in a 2mg/mL dopamine hydrochloride (Sigma Aldrich, Germany) solution in 10 mM Tris-HCl buffer (pH 8.5) under mild shaking for 4 hrs, as described earlier [7]. Subsequently, polydopamine-coated nanofibrous membranes were immersed in 3 mg/mL gelatin type B (Sigma Aldrich, Germany) solution in 10 mM Tris-HCl buffers, pH 8.5, overnight under shaking. Then, the gelatin-immobilized membranes were washed various times to remove residual gelatine from the fibrous mesh and dried at room temperature.

#### Evaluation of polydopamine coating and gelatin immobilization

We performed the microBCA assay (Thermo Fisher Scientific) to quantify the polydopamine coating on the nanofibers, following the previously reported method [8]. Membrane samples were cut into 0.5 cm² squares and were incubated with 200 μl of microBCA working solution for 2 hours at 37 °C. The reacted working solution was then measured by adsorption at 562 nm using Tecan Spark® Multimode Microplate Reader (Tecan Trading AG, Switzerland) to determine the coating amount. Parallel to this, standards were recorded, and the values were plotted against a standard curve to accurately determine the amount of polydopamine coating. Similarly, the microBCA assay was employed to indirectly quantify gelatin immobilization by measuring the remaining gelatin in the supernatant [9]. Gelatin release from the membrane has been measured over a span of 1, 3, 6, 9, 12 and 24 hours by keeping the membranes in PBS at 37°C while lightly shaking; 50 µL of supernatant is collected and evaluated as stated above.

The surface chemistry of the membranes, including PLLA Nanofibers (PL-NFs), PLLA Dopamine-coated Nanofibers (PLDA-NFs), and PLLA Gelatin-modified Nanofibers (PLGel-NFs), was analyzed using Raman spectroscopy. Membrane samples (1 cm²) were placed under a DXR Raman Microscope (Thermo Scientific, USA), equipped with a 10x objective. The membranes were irradiated with a 780 nm laser at an intensity of 15 kW.

Raman spectra were recorded and processed using Omnia software to identify chemical composition and surface modifications.

#### Surface characterization of nanofibers

We performed scanning electron microscopy (SEM) on the nanofiber membranes to visualize their surface morphology with secondary electron images. The samples were sputter-coated with a 4 nm platinum layer using a Leica EM ACE600 sputter coater (Leica Microsystems GmbH, Wetzlar, Germany) and examined using a Zeiss Crossbeam 340 Field-Emission Electron Microscope (Zeiss Microscopy, Oberkochen, Germany) at an acceleration voltage of 2 kV. To further assess the 3D structure and membrane thickness, Gallium Focused Ion Beam (Ga-FIB) imaging was employed. To evaluate membrane topography and fiber morphology SEM images have been analyzed with Image J. Porosity of the electrospun mat has been calculated from ratio between membrane mass and bulk PLLA as previously shown [6].

Additionally, atomic force microscopy (AFM) was conducted under ambient conditions using a Bruker Multimode 8 SPM system in tapping mode. Silicon cantilevers (OMCL-AC160TS, Olympus) with a resonance frequency of ∼300 kHz and a spring constant of ∼26 Nm ¹ were used. Carbon adhesive pads were utilized to fix the samples, including gelatin, PLLA, and PDA-coated membranes, onto magnetic metal discs for surface roughness and topography analysis.

#### Design and fabrication of a 3D-printed bioreactor for bilayer culture

To support bilayer culture in static conditions, we designed insert rings that facilitate the culture process. These rings were 3D printed using Fotodent® biocompatible resin (FotoDent® guide 385 and 405 nm, Dreve, Germany) with a DLP PRUSA SL1S speed printer. The membranes are securely fixed between two 3D printed rings.

For the bioreactor, we developed a millifluidic design using Computer-Aided Design (CAD) software, OnShape®. The bioreactor features a central channel measuring 1 mm in width, 1 mm in height, and 10 mm in length, with side wings designed for tubing connections. The top of the channel is open to allow for the insertion of a Polydimethylsiloxane (PDMS) membrane, which facilitates oxygenation. This system is also 3D printed using the DLP Prusa printer (Prusa SLS1 speed, Joseph Prusa Research, Czech Republic) with Fotodent® resin at a layer height of 0.025 µm.

Sylgard PDMS is prepared according to the manufacturer’s specifications, cast to form a 0.5 mm thick membrane, and subsequently cured. The PDMS membrane serves as the bioreactor’s window and is sealed using a biocompatible silicone (Dublisil, Dreve Dentamid GmbH, Unna, Germany) to ensure water tightness. After the membrane is securely placed between the two halves of the bioreactor, it is tightened using screw-driven pinchers (A. Hartstein, Germany) and connected to perfusion tubing via Luer lock. Two syringe pumps (brand to be specified) filled with media are connected to opposite sides of the bioreactor and programmed for perfusion. To verify laminar flow and shear stress on the membrane, we utilized the COMSOL Multiphysics software for laminar flow by setting as boundary condition the walls, inlet and outlet as per experimental values.

#### Cell culture of human immortalized podocytes and glomerular endothelial cells

Conditionally immortalized human podocyte cell line has been developed by Moin Saleem, Children’s and Renal Unit and Bristol Renal, University of Bristol, by retroviral transfection of a nephroctomy specimen with the temperature-sensitive *SV40-T* gene [10]. Therefore, cells can be expanded under permissive conditions at 33 °C. When cultivated at 37 °C, the transgene gets inactivated, cells enter growth arrest and undergo terminal differentiation. Human podocytes were cultured in RPMI Medium 1640 (Gibco, ThermoFisher Scientific, Waltham, MA, USA) supplemented with 10% heat-inactivated fetal bovine serum (FBS, PAN-Biotech, Aidenbach, Germany), 1%penicillin-streptomycin (Sigma-Aldrich, Merck, St. Louis, MO, USA), and 0.1% insulin-transferrin-selenium (ThermoFisher Scientific, Waltham, MA, USA).

Primary human glomerular microvascular endothelial cells (glomerular endothelial cells ACBRI 128) were purchased from Cell Systems, Kirkland, WA, USA, and were maintained in commercially available endothelial cell media (VascuLife® VEGF-Mv Medium, LifeLine® Cell Technology, ThermoFisher Scientific,Waltham, MA, USA) containing 5 ng/mL rhFGF basic, 50 g/mL ascorbic acid, 1 g/mL hydrocortisone hemisuccinate, 10 mM L-glutamine, 15 ng/mL rhIGF-1, 5 ng/mL rhEGF, 5 ng/mL rhVEGF, 0.75 U/mL heparin sulfate, 5% fetal bovine serum, 30 mg/mL gentamycin, and 15 g/mL amphotericin B (all supplements, LifeLine® Cell Technology, Thermo Fisher Scientific, Waltham, MA, USA).

#### hiPSC-culture and differentiation

HiPSC lines were produced by *Rose et al.* [11]. As described before. Human material was obtained from skin biopsies of healthy volunteers who gave written informed consent. Shortly, dermal fibroblasts outgrown from skin biopsies were episomally reprogrammed with a plasmid mix (pCXLE-hOCT3/4, pCXLE-hSK, pCXLE hMLN) using electroporation. For expansion and maintenance, hiPSCs were cultivated on a matrigel-coated (Corning, 354277, at a final concentration of 2.5 µg/ml) Nunc™ 6 well plate with mTeSR1 (Stem Cell Technologies, 85850) culture medium supplemented with Pen/Strep (Sigma-Aldrich, P4333). The medium was replaced every second day. Cells were passaged once or twice a week by treatment with accutase (Gibco, A1110501). To enhance cell survival, the accutase and the cell medium were supplemented with 10 µM Rho/associated kinase (Y-27623, Tocris, 1254) inhibitor for 24 h.

HiPS-derived podocytes were generated using previously described chemically defined growth medium conditions [4b, 4i, 12]. The differentiation protocol is summarized in Fig. 3A. First, cells were seeded in high density (1×10^4^ hiPSC/cm^2^) onto Nunc™ EasYFlask™ T75 cell culture flasks coated with Matrigel® as described above. After attaching overnight, the prewarmed human mesoderm differentiation medium (DMEM/F12 (Gibco^TM^, 11320074) supplemented with 1x B27 supplement (50x) (Gibco^TM^, 17504044), 1% Pen/Strep, 3 µM CHIRR99021 (Sigma-Aldrich, 252917-06-9), 50 ng/mL Activin A (Stem cell technologies, 78001.1), 10 µM Y27632) was added to the cell culture. After two days, it was replaced with the intermediate mesoderm differentiation medium (same as mesoderm differentiation medium except for the absence of Y27632 and an additional 50 ng/mL BMP7 (Peprotech, 120-03P)). The medium was changed every second day for the next 14 days. For the terminal differentiation into hiPSC-derived podocytes, cells were cultured in the podocyte end-differentiation medium (intermediate mesoderm differentiation medium with the addition of 25 ng/mL VEGF (Thermo Scientific, PHC9394) and 0.5 µM All-trans retinoid acid (Stem Cell Technologies, 72262). After differentiation, hiPSC-derived podocytes were maintained in the podocyte cell medium as described above.

#### Ethical statement

All preparations were approved by the local ethics committee (Approval no. 251_18B, Universitätsklinikum Erlangen). All experiments were performed in accordance with guidelines and regulations of the ethic committee of Universitätsklinikum Erlangen. Confirms that informed consent was obtained from all participants.

#### Artificial GFB characterization in static culture conditions

For artificial GFB under static conditions, PLLA membranes were fixed between two supporting rings and sterilized using a UV lamp. Suspension with 200.000 glomerular endothelial cells was seeded into the inner part of supporting rings (radius 3 mm) onto the membrane. The prewarmed human endothelial medium was added outside the supporting rings into the well to the brim of the supporting rings. The next day, the construction was turned upside down. 7×10^5^/cm^2^ pre-differentiated human imPodocytes or 3,5×10^5^/cm^2^ hiPSC-derived podocytes were seeded onto the membrane. The well was filled with a mix of human endothelial and podocyte medium to the brim of supporting rings. The day after, the constructs were placed into a 6-well plate containing 8 ml of mixed medium. On day four, the artificial GFB was characterized using the methods described below.

#### Artificial GFB characterization in dynamic culture conditions with the bioreactor

After sterilization with ethanol and /or UV light, a membrane was fixed between two parts of the bioreactor. Cell suspension containing 1×10^6^/cm^2^ glomerular endothelial cells were seeded into one chamber. Next day, glomerular endothelial cells were supplied with fresh medium and cell suspension of 1×10^6^/cm^2^ imPodocytes or 0.5×10^6^/cm^2^ hiPSC-derived podocytes were added into the second chamber. The day after, the bioreactor was connected to syringe pumps via tubing. Low flow rates (0.5 µl/min for podocytes, 1 µl/min for glomerular endothelial cell) were applied in the bioreactor for 24 h followed by four days of higher flow rates (5 µl/min for both cells).

#### Live dead assay

Cell viability was addressed using the LIVE/DEAD™ Cell Imaging Kit (ThermoFisher Scientific, Waltham, MA, USA). The kit was used according to the manufacturer’s instructions. Shortly, two components were mixed and added to samples (1:1 with cell medium). After the incubation of 15 min, fluorescence was visualized using EVOS™ M3000 Imaging System (ThermoFisher Scientific, Waltham, MA, USA).

#### Scanning Electron Microscopy analysis of artificial GFB

The cells grown on the membrane, referred to as GFB samples, were fixed using a solution of 4.5% paraformaldehyde (PFA) and 2% glutaraldehyde in 0.1 M HEPES buffer (pH 7.4). The samples were then dehydrated through a graded ethanol series and dried overnight with hexamethyldisilazane (HMDS). Following this preparation, the samples were sputter-coated, and scanning electron microscopy (SEM) was performed as previously described.

#### Immunofluorescent staining

*S*amples were washed with 1×PBS and fixed with 4% PFA (Roth,Karlsruhe,Germany) for 15 min. After rinsing, samples were incubated with blocking solution (1× PBS containing 10% normal goat serum (NGS, Abcam, Cambridge, UK), 1% bovine serum albumin (BSA, Karlsruhe, Germany), 0.5% Triton X–100 (Merck, Darmstadt, Germany)) for 1 h at room temperature (RT). Subsequently, samples were incubated with primary antibodies (supplementary table S1) diluted in 1× PBS containing 3% NGS and 1% BSA overnight at 4 ◦C. The next day, samples were washed three times with 1× PBS and labeled with fluorescent secondary antibodies (A-31634, goat anti rabbit 647 (Thermo Fisher Scientific, Waltham, MA, USA), goat anti-mouse 488 (A-21244, Thermo Fisher Scientific, Waltham, MA, USA) in the dark and at RT for 1 h. To label actin cytoskeleton, Alexa Fluor 555 Phalloidin (A-34055, Thermo Fisher Scientific, Waltham, MA, USA) was applied together with secondary antibodies.

Next, nuclei were stained with DAPI (Thermo Fisher Scientific, Waltham, MA, USA; 1:200 dilution in 1× PBS for 15 min in the dark). After washing with 1× PBS, samples were mounted with a drop of Fluoromount-G™ Mounting Medium (Invitrogen, Thermo Fisher Scientific, Waltham, MA, USA) between a cover and a microscope glass slide.

#### VEGF-A electroporation

A human VEGFA sequence coupled with a GFP sequence was cloned into a mammalian gene-expression vector (pLV[Exp]-Puro-CMV>3xFLAG/ORF_Stuffer, #VB900138-3673 twt, Vectorbuilder, Chicago, IL, USA). After transformation and expansion in *E. coli*, the plasmid was purified using the QIAprep Spin Miniprep Kit (QIAGEN). To transfect the VEGFA-GFP reporter plasmid terminally differentiated hiPSC-derived podocytes were prepared by washing with PBS and Opti-MEM Medium (ThermoFisher Scientific). Then, 5 µg of plasmid per 1–2.5 million cells were electroporated at 280V, Discharge Interval 50, 151MSEC (Progenetor 11, Hoefer). Cells were seeded onto the aligned membrane colonized with glomerular endothelial cells from the opposite side the day before. After three days of co-culturing, the artificial GFBs were fixed and mounted as described above and visualized using confocal microscopy.

Undifferentiated conditionally immortalized human podocytes were transfected with 5 µg VEGFA plasmid using the ProGenetor II electroporator (Hoefer, Holliston, MA, USA) with one pulse at 280 V and 1200 µF.

#### MiRNA transfection

After seeding and attachment on the aligned membrane, glomerular endothelial cells were transfected with 200 nM Alexa555-tagged miR-192 (MirVanaTM, custom miRNA mimic, Thermo-Fisher Scientific) for 4 hours using Lipofectamine 2000 and Opti-MEM Medium (ThermoFisher Scientific) according to manufacturer’s protocol. The next day terminally differentiated hiPSC-derived podocytes were labeled with eBioscience Cell Proliferation Dye eFluor 450 (Invitrogen, final concentration of 2 µM) according to the manufacturer’s instruction and seeded onto the opposite side of the membrane. After 24 h of co-culturing, the artificial GFBs were fixed, nuclei were stained with SYTO Deep Red (Thermo-Fisher Scientific) for 5 min, rinsed, mounted as described above and visualized using confocal microscopy.

#### Permeability assay

Artificial GFBs without cells or seeded with glomerular endothelial cells and imPodocytes were moved to a clean well. A dextran mix consisting of Cascade Blue-labeled 10 kDa and Texas Red-labeled 70 kDa dextrans (Thermo-Fisher Scientific) diluted in human endothelial and podocyte medium mix (100 µg/ml, 50 µl) was added into the inner part of supporting rings (radius 2 mm). Pure human endothelial and podocyte medium mix was added outside the supporting rings into the well. After 30 min of incubation (37°C), the artificial GFB construct was moved out and the remaining liquid was collected. The fluorescence was measured using GloMax-Multi+Detection System (Promega) at 405/495 and 625/660 excitation/emission wavelength, respectively.

### Statistical Analysis

Data is presented as mean ± standard deviation and the replicates per condition are at least *n* = 3 unless declared otherwise. Statistical evaluation was performed using GraphPad Prism (version 9.5.1) or MATLAB(R) (version R2022a) using unpaired t-test. Statistically significant differences between groups are indicated by * (*p* <0.05) or **p<0.01.

## RESULTS

### Characterization of the functionalized electrospun nanofibers

We used electrospinning to produce an artificial GBM that was later seeded with human glomerular endothelial cells and human immortalized or hiPSC-derived podocytes on opposite sides of this membrane. The electrospun nanofibers offer numerous advantages over the commercially available synthetic systems in terms of size, shape, variety of materials, fiber morphology, and orientations, and the modification of nanofibers with bioactive cues [6]. Poly-L-Lactic Acid (PLLA) was used for electrospinning due to its stronger mechanical properties that allowed us to achieve fibers in nano-/micrometer size and to gain enough mechanical stability for handling. The polymer was electrospun in a random or aligned topography, and cell behavior was compared to find optimal conditions for our purposes (Fig. 1 A and B). The electrospun mats presented an average thickness of 7 µm, alignment was present in 86% of the fibers (Fig. 1C), and an average fiber diameter of 0.6±0.4 µm (Fig. 1D). The porosity of the mat was calculated at about 57% and 63% for random and aligned, respectively.

**Figure 1:**
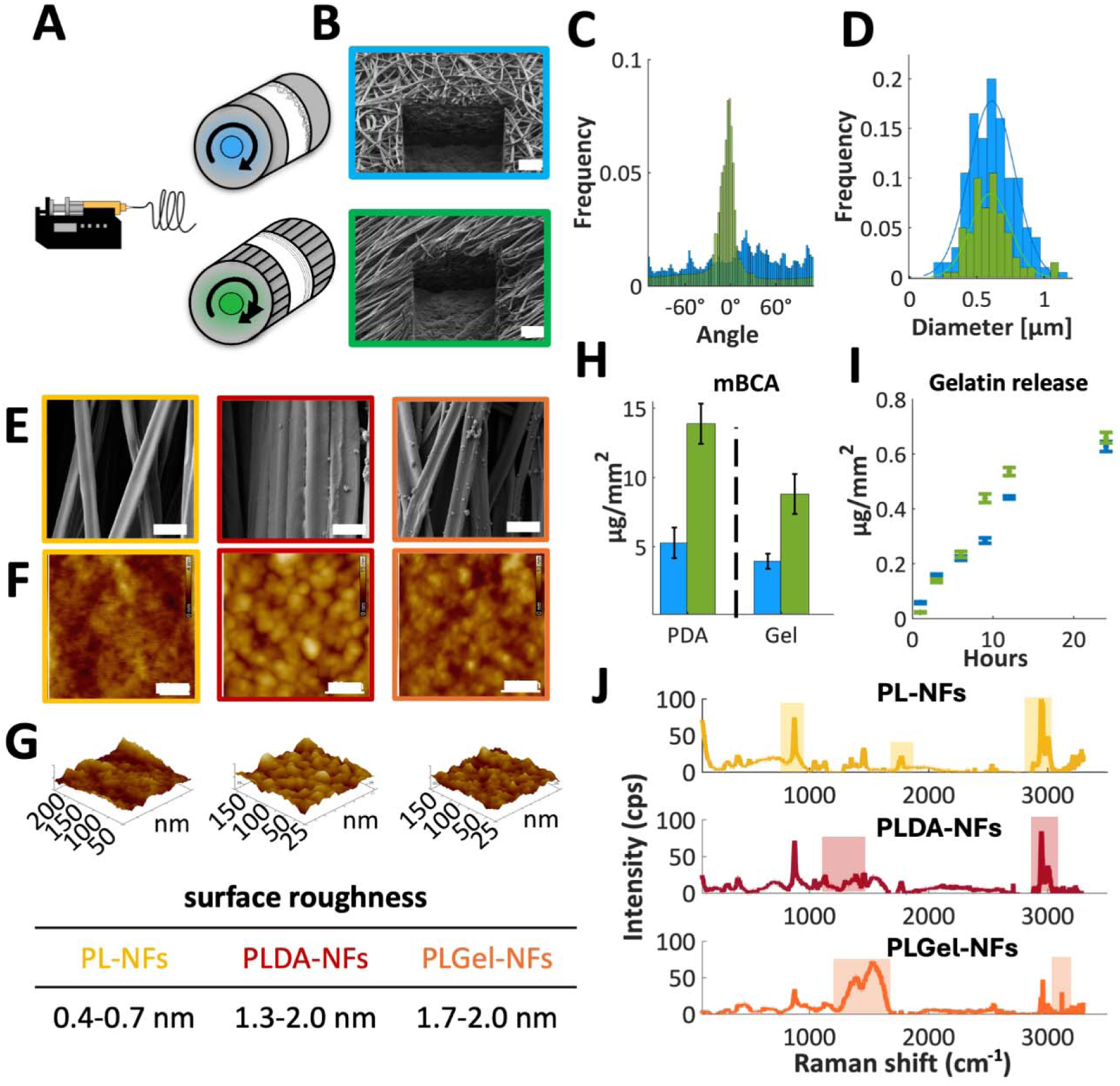
Characterization of nanofiber-based artificial membrane. (A) Schematic illustration of electrospinning random and aligned membranes using different mandrels. B: SEM images with Gallium focused-ion-beam of random (blue) and aligned (green) nanofibers. Scale bar 4 µm. (C) Fiber alignment distribution for random (blue) and aligned (green) membranes. (D) Fiber diameter distribution for random (blue) and aligned (green) mats. (E) SEM images of PL-NF, polydopamine PLDA-NF and gelatin PLGel-NF coated fibers. Scale bar 1 µm. (F) AFM Image of PL-NF, polydopamine PLDA-NF and gelatin PLGel-NF. Scale bar: 50 nm. (G) AFM surface roughness analysis (Root Mean Square). (H): mBCA quantification of coating for random (blue) and aligned (green) membranes. (I) Gelatin release from membrane over 24 hours. (J) Raman spectroscopy of surface chemistry for PLLA, PDA and gelatin. Abbreviations: AFM: atomic force microscope, PL-NF: Poly-L-lactic acid nanofibers, PLDA-NF: Polydopamine coated PLLA nano fibers, PLGel-NF: Gelatin coated PLLA nano fibers, SEM: Scanning electron microscopy.

We surface-modified the PLLA nanofibrous membranes by immobilizing gelatin via polydopamine chemistry. Scanning electron microscopy (SEM) imaging revealed no significant changes in the diameter of nanofibers before or after coatings (Fig. 1E). Interestingly, atomic force microscopy (AFM) measurements revealed a change in surface roughness (root mean square), 0.4-0.7 nm for PLLA, 1.3-2.0 nm for PDA and 1.7-2.0 nm, for gelatin surfaces (Fig. 1F and G). The quantification of the amounts of coating was performed with a micro-BCA assay. This analysis demonstrated about 3.96 ± 0.55 µg of gelatin for the random topography and 8.82 ± 1.45 was coated per mm^2^ of nanofiber (Fig. 1H). Measurement of gelatin release over time showed that up to 0.6 µg/mm^2^ of gelatin detached from the mat in 24 hours (Fig. 1I).

To prove the technique’s efficacy, we analyzed the fibers’ surface by Raman spectroscopy. Characteristic Raman peaks could be detected for the chemical bond molecules and confirmed the property of the surface chemistry (Fig. 1J). PLLA typically peaks at 873 cm^−1^ C-COO that can be attributed to the 1750 cm^−1^ identifies the C=O and at 2950 cm^−1^ to the CH_3_ chains. The dopamine spectrum differs for the presence of slightly higher peaks at 1500 and 1350 cm^−1^ suggestive of aliphatic and aromatic groups. Furthermore, the amine bond can be detected in the band of 3150 cm^−1^. The gelatin spectrum is mostly associated with the presence of ample peaks around 1560, 1650 and 3100 cm^−1^ due to amide bonds [13].

### Human glomerular cells on the artificial GBM

After these basic characterizations, the electrospun membrane was used for cell culture. Different numbers of human glomerular endothelial cells and differentiated immortalized podocytes were seeded onto random or aligned directed electrospun PLLA fibers as monocultures to find cell numbers and conditions best for each cell type. The fibers were either uncoated or functionalized with gelatin Type B using polydopamine chemistry to improve cell adhesions. To establish a tri-layered model for the GFB with an electrospun PLLA membrane as support for the co-culture of glomerular cells, the membrane was fixed between two 3D printed supporting rings allowing to seed glomerular endothelial cells and podocytes on different sides of the membrane (Fig. 2A, B). By fabricating the artificial GBM in different topography, we demonstrated that cell attachment and morphology were impacted by fiber topography and co-culture. In co-culture with glomerular endothelial cells on the other side of the PLLA membrane podocytes survived on both fiber topographies (shown by life dead staining). However, immortalized podocytes cultured on a random membrane formed aggregates and the layer exhibited considerable gaps between cells. On the other hand, podocytes cultured on an aligned membrane demonstrated tight monolayer and elongated morphology shown by immunofluorescent staining and SEM (Fig. 2C). Likewise, glomerular endothelial cells only formed a coherent monolayer on aligned membrane but not on random membrane (Fig. 2D).

**Figure 2:**
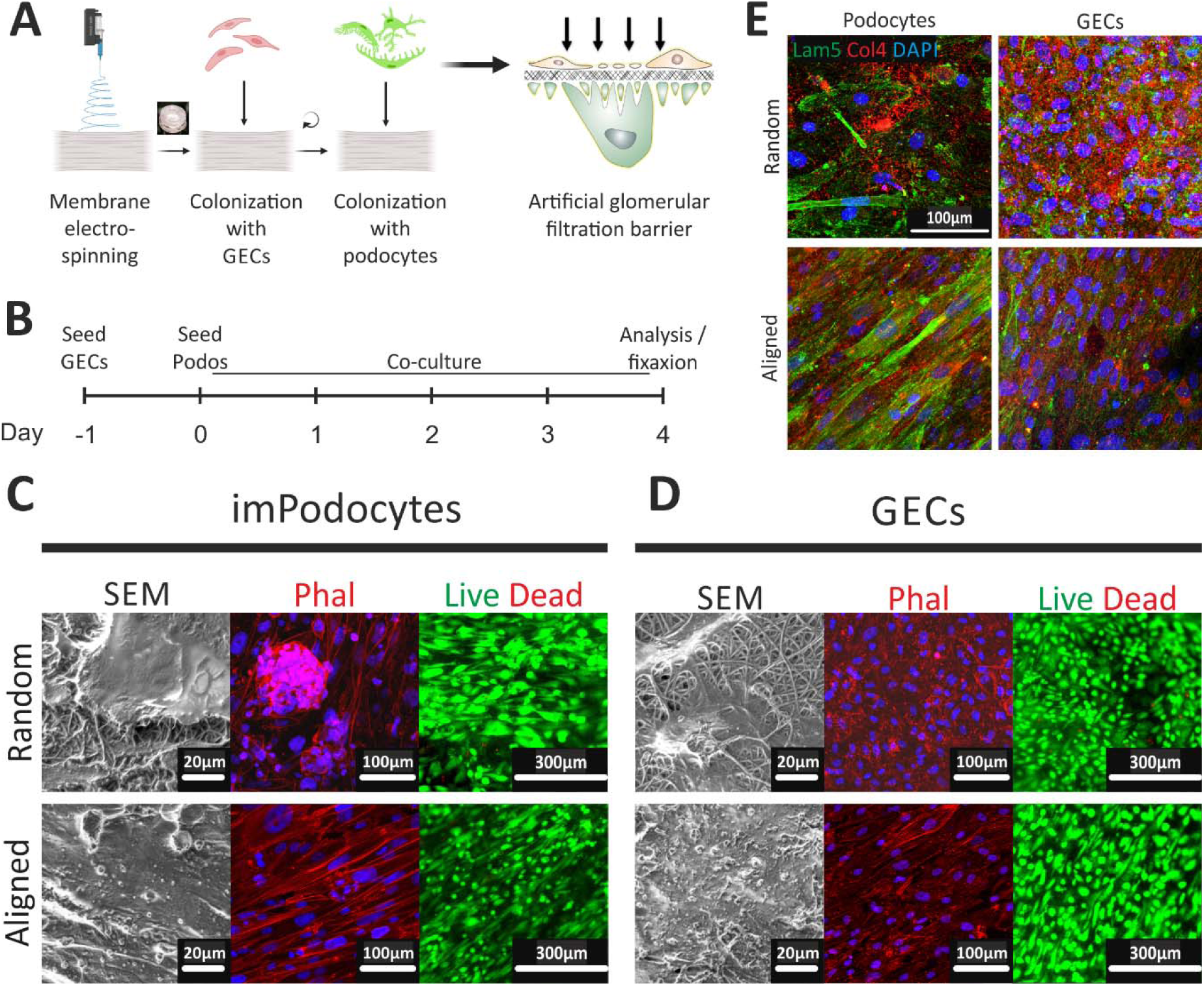
Co-culture of human conditionally immortalized podocytes (imPodocytes) and glomerular endothelial cells (GECs) on the electrospun polydopamine- and gelatin-coated PLLA-membrane. (A) Schematic protocol for membrane colonization with human GECs and imPodocytes from different sides of the membrane. Created in BioRender. Laptii, A. (2023) BioRender.com/g63h412. (B) Schematic timeline of cell co-culturing on the electrospun membrane. (C-D) Layer integrity and cell morphology of (C) imPodocytes and (D) GECs on the membrane with random (upper panel) and aligned (bottom panel) topography visualized using SEM (left panel), confocal microscopy with phalloidin (Phal) staining (middle panel), and live/dead staining (right panel). Scale bar 20 µm, 100 mm, and 300 mm respectively. (E) Immunofluorescent staining for laminin 5 (Lam5; green), collagen IV (Col4; red), and DAPI nuclei staining (blue) of imPodocytes and GECs cultured on the membrane with random (top) or aligned (bottom) topography. Scale bar 100 µm. Abbreviations: GECs: glomerular endothelial cells, ImPodocytes: human conditionally immortalized podocytes, SEM: Scanning electron microscopy.

Glomerular endothelial cells and podocytes are known to produce extracellular matrix (ECM) of the GBM. We could show that important components of the natural GBM such as laminin-521 and collagen α3α4α5 (IV) were synthesized by our cells themselves within our model, especially when cells were cultured on aligned fibers (Fig. 2E).

### hiPSC-derived podocytes show better podocyte characteristics compared to imPodocytes

Immortalized podocytes have several limitations: Cells can dedifferentiate in culture, especially when they reach confluency, and several podocyte-specific markers are either only slightly or not expressed at all [14]. This brings the use of immortalized podocytes and their applicability for physiological, pathophysiological, and clinical research into question. Recently, we established a protocol for the generation of human podocytes - including patient-specific podocytes - from a skin punch biopsy by episomal reprogramming of dermal fibroblasts into human-induced pluripotent stem cells (hiPSCs) and subsequent differentiation into hiPSC-derived podocytes (Fig. 3A) [12]. We introduced hiPSC-derived podocytes into our *in vitro* model of the GFB. HiPSC-derived podocytes demonstrated high detachment from the membrane cultured as a monoculture (data not shown). In contrast, hiPSC-derived podocytes cultured in combination with glomerular endothelial cells on the opposite side of the membrane displayed a high viability and tight monolayer, at least for four days of culturing (Fig. 3B). Indeed, podocytes generated by our protocol resemble *in vivo* podocytes much better in terms of morphological characteristics and expression of podocyte-specific markers. Again, hiPSC-derived podocytes survived on aligned and random fibers but culturing cells on aligned fibers allowed for higher cell confluency and better monolayer formation of hiPSC-derived podocytes (Fig. 3B) and glomerular endothelial cells (Fig. 3C).

**Figure 3:**
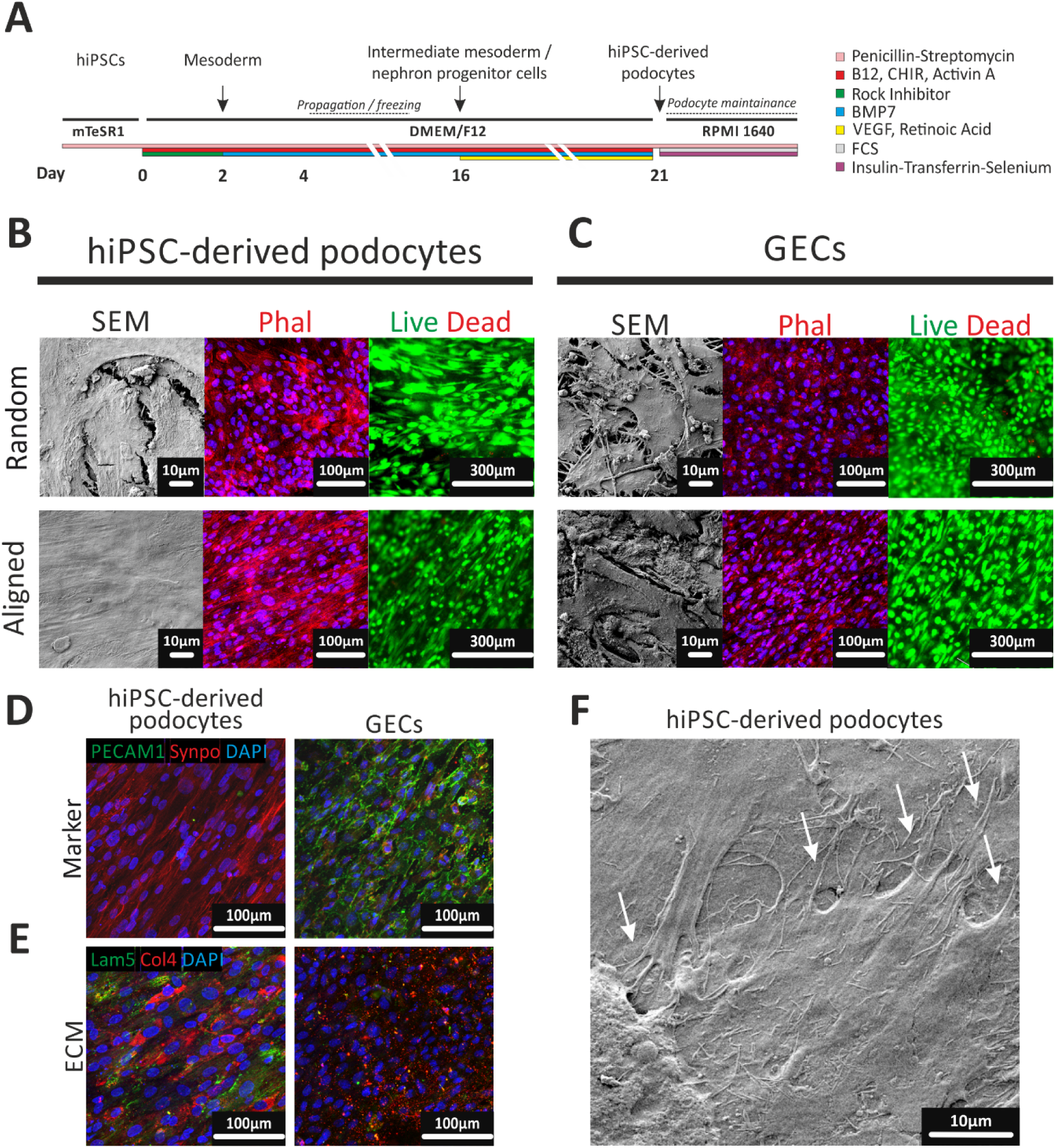
Co-culture of hiPSC-derived podocytes and GECs on the electrospun polydopamine- and gelatin-coated PLLA-membrane. (A) Schematic timeline and medium composition for hiPSCs-differentiation into podocytes. (B-C) Layer integrity and cell morphology of (B) hiPSC-derived podocytes and (C) GECs on the membrane with random (top) and aligned (bottom) topography visualized using SEM (left), confocal microscopy with Phal staining (middle), and live/dead staining (right). (D-E) Expression of the (D) cell-type specific marker and (E) ECM in the co-culture of hiPSC-derived podocytes (left) and GECs (right) cultured on the opposite sides of the membrane with aligned topography. Membranes were stained against (D) PECAM1 (green) and synaptopodin (Synpo, red) as well as (E) laminin 5 (Lam5, green) and collagen type IV (Col4, red) and visualized using confocal microscopy. (F): Podocyte-like cell protrusions (white arrows) of the hiPSC-derived podocytes co-cultured with GECs on the membrane with aligned topography visualized using SEM. ECM: extracellular matrix, GECs: glomerular endothelial cells, hiPSC: human induced pluripotent stem cells, PECAM1: Platelet and endothelial cell adhesion molecule 1, SEM: Scanning electron microscopy.

Podocyte-specific marker synaptopodin was expressed on the hiPSC-derived podocyte side of the membrane whereas the endothelial cell marker PECAM1 was found as a zipper-like structure along the cell edges on the other side of the membrane (Fig. 3D). Collagen IV was expressed by both cell types whereas laminin 5 was only secreted by hiPSC-derived podocytes (Fig. 3E). Moreover, hiPSC-derived podocytes showed typical foot-like processes between neighboring cells (Fig. 3F).

### Cell communication through the artificial GFB

Cell-cell communication is important for proper functioning of the GFB. Both, glomerular endothelial cell to podocyte and podocyte to glomerular endothelial cell communication happens through the natural GBM [15]. It is known that glomerular endothelial cells need VEGF derived from podocytes for formation of fenestrae [16]. We electroporated human podocytes with a plasmid encoding fluorescent VEGF-A and co-cultured the cells with glomerular endothelial cells on the PLLA membrane. Imaging the membrane from the endothelial side could confirm uptake of podocyte-derived VEGF-A by glomerular endothelial cells (Fig. 4A).

**Figure 4:**
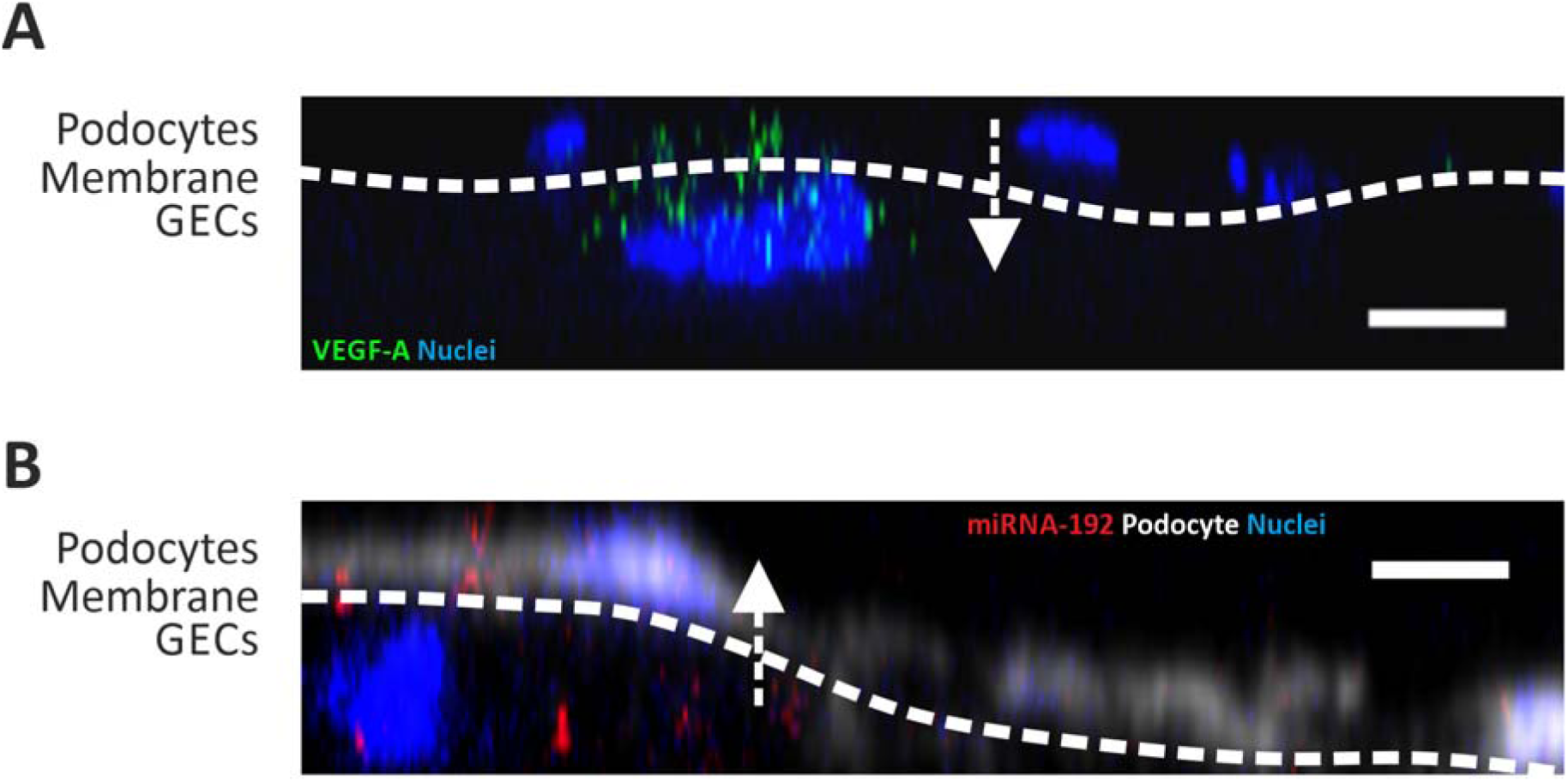
Cell communication through the electrospun membrane. (A) Podocyte-secreted VEGF-A (green) crossed the electrospun membrane and was taken up by GECs. (B) Fluorescent-labeled miRNA-192 (red) secreted from GECs was detected within hiPSC-derived podocytes labeled with eFluor450 (white). Visualized using confocal microscopy from the GEC- (A) or podocyte-side, white arrows show the direction of signaling, scale bar 10 µm. Abbreviations: GECs: glomerular endothelial cells, hiPSC: human induced pluripotent stem cells.

We were also able to show cell-cell communication in the opposite direction. It is known that glomerular endothelial cells and podocyte communicate through microRNAs (miRs) [17]. We transfected glomerular endothelial cells with a red fluorescent miR-192 and co-cultured the cells with podocytes on the artificial GBM. We could detect the uptake of the red fluorescent endothelial cell-derived miR in podocytes (Fig. 4B).

Thus, we could show that cell-cell communication between glomerular endothelial cells and podocytes was possible in both directions.

### Artificial GFB shows size selective permeability

Different molecular weights of fluorescent dextrans were added to the endothelial site of the artificial GFB fixed between two rings under static conditions. Aligned fibers alone without any cells allowed the passage of 10 kDa as well as 70 kDa dextrans (Fig. 5A-C). After seeding glomerular endothelial cells and podocytes on opposite sides of the artificial GBM 10 kDa dextran was still able to pass the artificial GFB but in much lower amount. After 30 min around 2% of 10 kDa dextran added to the endothelial side was detectable on the podocyte side of the membrane (Fig. 5A). In contrast, no 70 kDA dextran passed the artificial GBM seeded with glomerular cells after 30 min indicating its tightness (Fig. 5B, C). Thus, our *in vitro* GFB shows permselectivity comparable to *in vivo* conditions.

**Figure 5:**
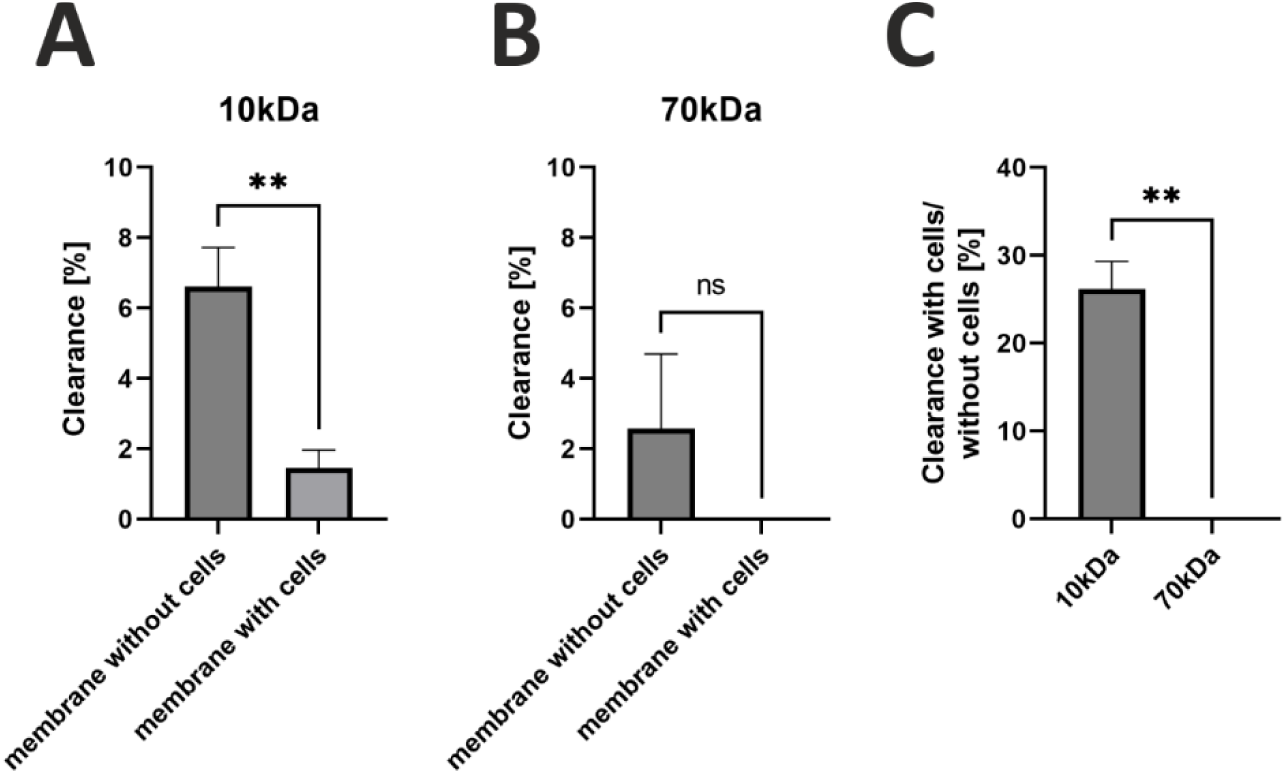
Permeability of the AFB. (A) Percentage of clearance of 10 kDa dextran (added to the GEC-side) through the AFB after 30 min. **p<0.01. (B) Percentage of clearance of 70 kDa dextran (added to the GEC-side) through the AFB after 30 min. (C) Comparison of clearance of 10kDa and 70kDa with podocytes and GECs on the membrane versus membrane without cells. **p<0.01. Unpaired t-test, n=3. Abbreviations: AFB: artificial filtration barrier, GECs: glomerular endothelial cells.

### Flow-induced shear stress caused cell maturation with fenestrae development

Next, we implemented flow into our model by incorporating the membrane in a bioreactor with the intention to improve endothelial cell maturation by shear stress. The bioreactor was designed on OnShape CAD software and printed with a PRUSA SLS1 Direct Light Processing (DLP) printer in Fotodent(R), a biocompatible resin (Fig. 6A, B). Advantages of such bioreactors compared to classic organs-on-chip are in the possibility to design and print a stable custom-made device that can be sterilized and reused multiple times while still achieving the sub millimeter resolution of DLP Printers.

**Figure 6:**
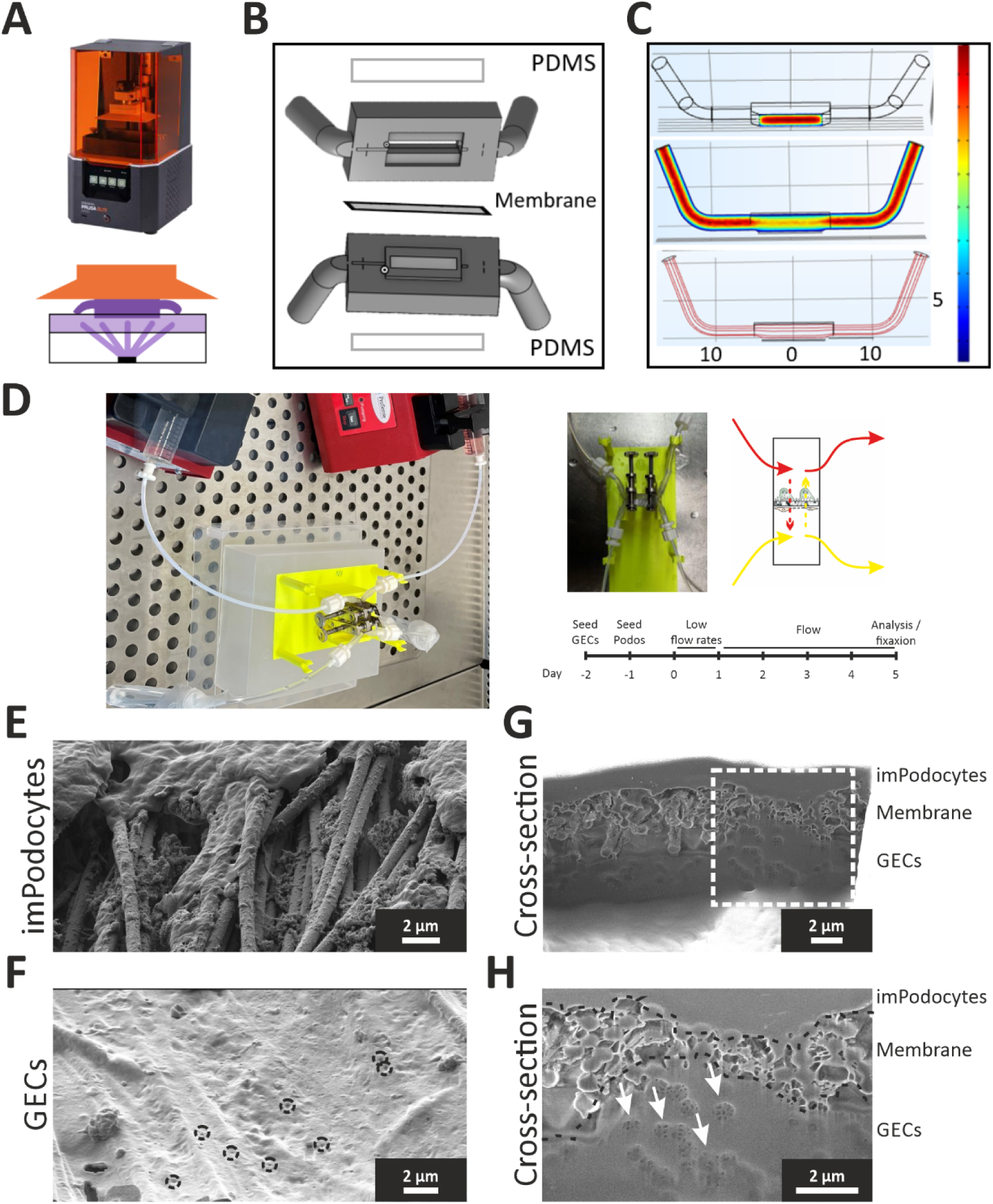
Design of the bioreactor and flow effects. (A) Direct light processing 3D printing of the bioreactor with a Prusa SL1S printer (B) Schematic of bioreactor components. (B) Comsol simulation of flow: shear stress scale bar 0-18 mPa (top), flow velocity scale bar 0-12 mm/s (middle), flow lines (bottom). (D) Bioreactor perfusion setup and experimental timeline. (E-F) Detection of foot process-like (E) and fenestrae-like (F, black circles) structures by imPodocytes and GECs, respectively, using SEM. (G-H) A cross-section of the AFB with fenestrae-like structures organized in clusters (white arrows) using focused-ion-beam SEM. Abbreviations: AFB: artificial filtration barrier. GEC: glomerular endothelial cells, hiPSC: human induced pluripotent stem cells, imPodocytes: human conditionally immortalized podocytes, SEM: Scanning electron microscopy.

Since the resin is not transparent and does not transpire for oxygen, a pre-cast thin PDMS membrane was added to the reactor to allow oxygenation. The reactor sealed mechanically with the membrane fixed in between two parts compressed with pinchers. The bioreactor presents two inlets and two outlets to allow parallel flow on the endothelial side and on the podocyte side. A model of the flow in the bioreactor was simulated in the software “Comsol Multiphysics” to evaluate velocities and shear stress of the laminar flow that act on the cells on the membrane. Thereby, we aimed to optimize flow velocity and bioreactor design to better fit the physiological conditions (Fig. 6C).

Glomerular endothelial cells were seeded first. The next day, podocytes were seeded on the other side of the membrane. Flow was added to the model with the help of the bioreactor for 5 days (Fig. 6D). Due to the flow condition podocytes formed longer foot process-like structures around the fibers (Fig. 6E). Most importantly, induction of flow to our model induced the development of fenestrae-like structures in glomerular endothelial cells as an important hint for better maturation of glomerular endothelial cells in co-culture and under shear stress (Fig. 6G, H). Such degree of maturation of glomerular endothelial cell fenestrae was not show in other *ex vivo* models before.

## DISCUSSION

The GFB consists of three layers. The first layer is built of specialized glomerular endothelial cells characterized by a high number of perforations called fenestrae. The second layer is formed by a GBM consisting mainly of collagen IV, laminins, fibronectins, and proteoglycans. The last layer is formed by podocytes with interdigitating foot processes that are bridged by slit diaphragms. All layers of the GFB are important for its specific permselectivity [18]. There is an unmet need for *in vitro* GFB models to simulate physiological and pathophysiological conditions.

Most existing *in vitro* models for the GFB used commercially available on-chip devices [4a-i, 4k, 4l], tissue culture inserts or membranes [4g, 5] allowing to seed cells on opposite sides of a plastic membrane only in predefined conditions. For example, in the model by *Li et al*. 2016 commercially available and prescribed 1 μm porous membranes made of hydrophilic polytetrafluoroethylene (PTFE), polycarbonate or polyethylene terephthalate (PET) were used [5a, 5b]. However, OOC systems rely on synthetic membranes which are significantly thicker than human GBM and lack the necessary topographical features to support proper ECM remodeling and transmembrane crosstalk. Furthermore, most glomerular co-culture models of the past used murine cell lines [4g, 4k, 4l, 5b] or human immortalized podocytes [4c, 5a, 19] that do not resample *in vivo* podocytes in terms of podocyte maker expression and foot processes [14]. *Musah et al.* established a differentiation protocol of hiPSCs to vascular endothelium and podocytes and co-cultured them in a microfluidic kidney glomerulus chip for the first time [4i]. Although mechanical stimuli such as shear or tensile stresses are essential for the physiological functions of the GFB [20], only a few models of the past applied dynamic flow conditions [4e, 21]. For example, *Slater et al*. used human cells in their GFB model but did not apply flow and thus did not address filtration in their models [5a].

We used electrospinning to generate an artificial GBM that was later seeded with human glomerular endothelial cells and human immortalized podocytes but also stem cell-derived podocytes on opposite sides of this membrane. The surface of the nanofibrous membrane was either uncoated or functionalized with gelatin using polydopamine chemistry to improve cell adhesions and fibers were characterized by electron microscopy, Raman spectroscopy and FTIR spectroscopy. Different fiber orientations within the artificial membrane were tested and revealed that cell-cell and cell-material interactions as well as cell morphology were best maintained with cells seeded on aligned fibers. Thus, our surface functionalized artificial GBM supported a co-culture of glomerular cells to mimic the GFB.

Immortalized podocytes used by others as well as in our model have several limitations: Cells can dedifferentiate, and many podocyte-specific markers are not expressed. This brings the use of immortalized podocytes into question. However, we could show that co-culture with glomerular endothelial cells significantly improves morphology and marker expression of immortalized podocytes [22]. Furthermore, not only immortalized podocytes survived on the artificial GBM but also hiPSC-derived podocytes that revealed a morphology much more comparable to podocytes *in vivo*. HiPSC-derived podocytes were generated via a protocol starting form fibroblasts derived from a skin biopsy over electoporation of stem cell plasmids and final chemically defined differentiation into podocytes [12]. Glomerular cells produced their own extracellular matrix and expressed cell-type specific markers on the artificial GBM.

Furthermore, we were able to show that cell-cell communication between glomerular endothelial cells and podocytes was possible in both directions. This bidirectional communication is essential for cell maturation and function of the GFB as well as for glomerular integrity *in vivo* [15].

We assessed the permeability of the artificial GFB by measuring the passage of different-sized dextrans through the barrier. We could show that aligned fibers alone already formed a significant barrier for 10 kDa and 70 kDa dextrans. After seeding glomerular endothelial cells and podocytes the barrier was much tighter for 10 kDa dextran that was still able to pass. However, the artificial GFB seeded with glomerular cells did not allow the passage of 70kDa dextrans any longer. In vivo, the functional cut-off of the human GFB is also believed to be around 70 kDa and 10 kDa dextran injections in zebrafish showed similar clearance rates as our *in vitro* model [23]. Therefore, our artificial membrane revealed functionality that was very similar to *that of in vivo* conditions.

Shear stress that glomerular endothelial cells experience through blood flow and podocytes through urine flow within the Bowman capsule is important for cell maturation. We implemented a flow of 5 µl/min on the endothelial cell as well as on the podocyte side of our model by incorporating the membrane in a bioreactor. Indeed, we were able to show the flow-dependent formation of glomerular endothelial cell fenestrae that never developed in other glomerular endothelial cell models *ex vivo*. The podocyte foot process-like structures also matured after exposure to shear stress.

In summary, we could show that our artificial GFB enables *in vivo*-like transmembrane intercellular crosstalk, ECM production, cell maturation and perspectivity.

Compared to other 3D models of the GFB published before our model has the following advantages and novelties: Firstly, our artificial GBM allows flexibility in design concerning tunable porosity, control over fiber diameter, fiber orientation, and surface modification. Published models usually use commercially available chips that are not customized. Secondly, our surface modification methods allow for the functionalization of artificial GBM with bioactive cues. We are equipped with state-of-the-art instruments to characterize and validate the *in vitro* models allowing us to gain important information about the suitability of the model and the opportunity to tune it further depending on cell responses. Third, our functional assays, including the measurement permeability for different-sized dextrans as a sign of leakiness of the filtration barrier and the assessment of paracrine communication within the cells of the GFB, make the model suitable for later use regarding *ex vivo* studying of pathophysiological conditions and drug testing.

## ACKNOWLEDGMENTS

**General:** The authors would like to thank Sven Heilig for proofreading the manuscript and Victoria Rose for providing iPSCs. CM is supported by the GSLS Würzburg, Germany.

## Author contributions

CM and AR contributed equally to this work. CM: conceptualization, data collection, data analysis, methodology, visualization and writing. AR: conceptualization, data collection, data analysis, methodology, visualization and writing. PS: SEM imaging. VS, FW: AFM measurements. JG: funding acquisition, supervision. MS: Writing review, TA and JMD share correspondence and contribute equally to planning, project administration, supervision, funding acquisition, writing original draft, writing review editing.

## DATA AVAILABILITY

The datasets generated during and/or analyzed during the current study are available from the corresponding author on reasonable request

## SUPPLEMENTARY MATERIALS

**Supplementary table S1:** Primary antibodies.

| Antigen (Clone) | Host | Company, Cat. No. | Dilution |
| --- | --- | --- | --- |
| Collagen IV | rabbit | Ab6586, Abcam, Cambridge, UK | 1: 200 |
| Laminin 5 | mouse | Ab77175, Abcam, Cambridge, UK | 1: 200 |
| CD31 | mouse | Ab9498, Abcam, Cambridge, UK | 1: 200 |
| Synaptopodin | rabbit | 21064-1-AP, Proteintech, Rosemont, IL, USA | 1: 200 |

